# Re-modeling of foliar membrane lipids in a seagrass allows for growth in phosphorus deplete conditions

**DOI:** 10.1101/666651

**Authors:** Jeremy P. Koelmel, Justin E. Campbell, Joy Guingab-Cagmat, Laurel Meke, Timothy J. Garrett, Ulrich Stingl

## Abstract

We used liquid chromatography high-resolution tandem mass spectrometry to analyze the lipidome of turtlegrass (*Thalassia testudinum*) leaves with extremely high phosphorus content and extremely low phosphorus content. Most species of phospholipids were significantly down-regulated in phosphorus-deplete leaves, whereas diacylglyceryltrimethylhomoserine (DGTS), triglycerides (TG), galactolipid digalactosyldiacylglycerol (DGDG), certain species of glucuronosyldiacylglycerols (GlcADG), and certain species of sulfoquinovosyl diacylglycerol (SQDG) were significantly upregulated, explaining the change in phosphorus content as well as structural differences in leaves of plants growing under diverse phosphate concentrations. These data suggest that seagrasses are able to modify the phosphorus content in leaf membranes dependent upon environmental phosphorus availability.

## Main Text

Seagrasses are a widely distributed group of marine plants that provide a range of ecological services to coastal habitats around the world. Seagrass beds are in worldwide decline, mostly due to anthropogenic changes in nutrient delivery to coastal waters^1^. For over 30 years, it has been known that many seagrass species can display shifts in foliar phosphorus (P) content in response to environmental availability^2,3^, an adaptation that allows them to grow in a wide range of habitats with divergent nutrient conditions. In *Thalassia testudinum* (turtlegrass), a dominant species in South Florida,^4^ elemental C:P ratios can differ by more than 10 fold from around 200 to nearly 3000, often dependent upon environmental P availability.^3,5,6^ *Thalassia testudinum* is distributed along the western Atlantic from Florida, USA to Venezuela, throughout the Gulf of Mexico and the Caribbean Sea.^7^ *Thalassia hemprichii*, the other species in this genus, is also widely distributed in the coastal waters of the Indian Ocean and the western Pacific.^8^ While the morphology of turtlegrass leaves and canopy structure changes with decreased P content, areal production rates can remain relatively high,^6^ indicating metabolically active plants. Changes in C:P ratios and P content of turtlegrass occur along natural P gradients,^4,6,9^ but can also be induced by fertilization experiments in P-depleted habitats^5,10^. The exact cellular mechanisms on how turtlegrass lowers its P content are mostly unknown. In this study, we used liquid chromatography high-resolution tandem mass spectrometry (LC-HRMS/MS) to analyze the lipidome of turtlegrass leaves that contained either a high percentage of P (0.445 ± 0.017%) or a low percentage of P (0.083 ± 0.002%). The N, C, and P leaf content is shown in Table S1. The samples were taken from a fertilization experiment of P-limited turtlegrass in Largo Sound, FL, USA.^5^ In total, 600 unique molecular lipid species across 36 lipid classes (supplemental table: SeaGrassData_Supplemental.xlsx) were tentatively annotated by LipidMatch Flow^11,12^. The total lipidome of the samples grouped based on foliar P content without any exceptions (Figure 1).

**Figure 1:**
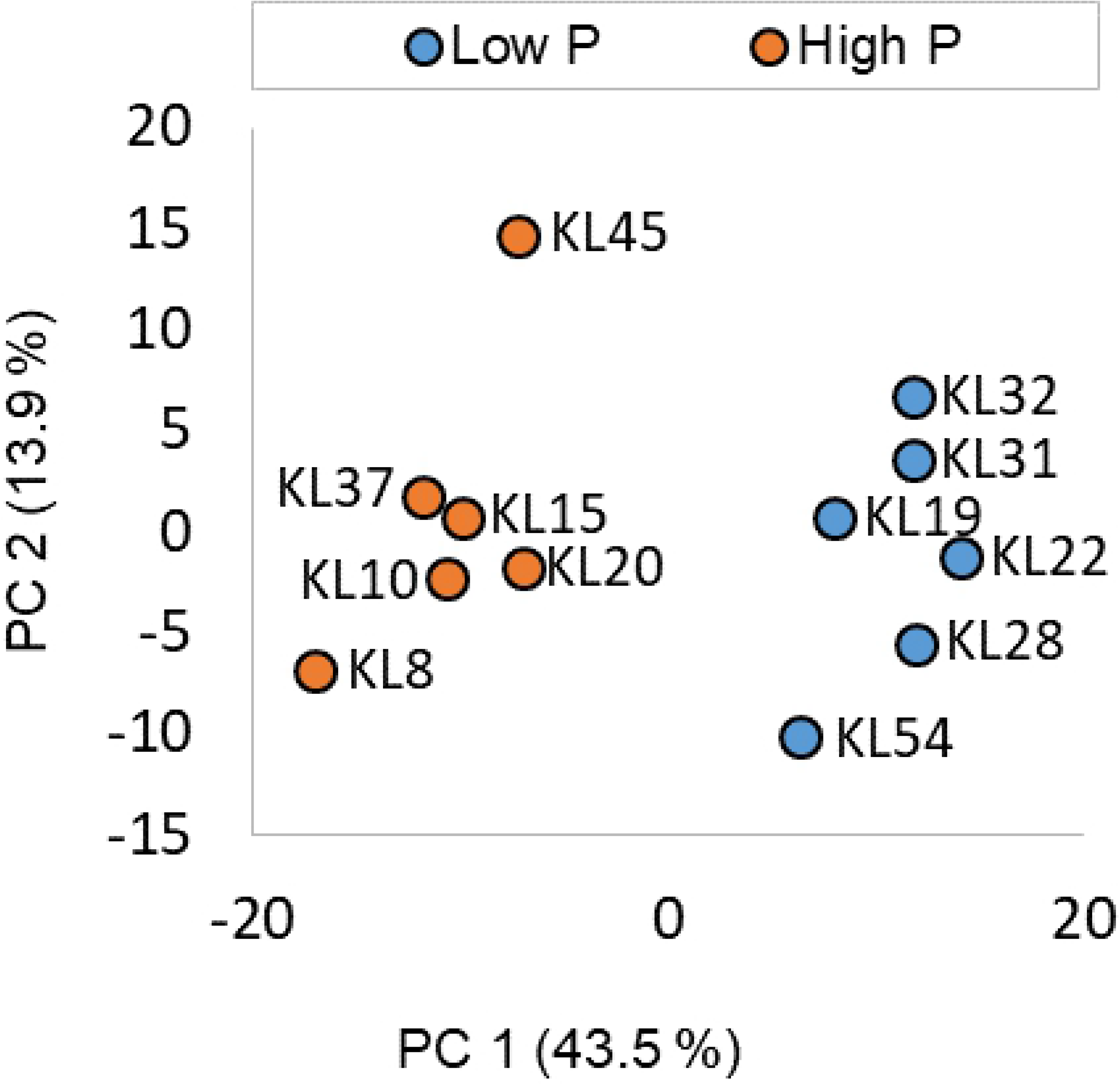
Principal Component Analysis of the total lipidome of samples with high phosphorous content (organge) versus samples with low phosphorous content (blue; normalized by sum, log transformed and mean centered). PC1 explained 43.5% of the variation, and PC2 explained 13.9% of the variation.

Most classes of phospholipids were significantly down-regulated in P-depleted leaves including PC and PE, which were reported as the most abundant phospholipids in three species of seagrasses,^13^ whereas diacylglyceryltrimethylhomoserine (DGTS), triglycerides (TG), galactolipid digalactosyldiacylglycerol (DGDG), certain species of glucuronosyldiacylglycerols (GlcADG), and certain species of sulfoquinovosyl diacylglycerol (SQDG) were significantly upregulated (Table 1, Figure S2) and presumably replace phospholipids in the membrane.

**Table 1:** Up- and down-regulated lipid classes based on a fisher’s exact test (see supplementary methods for details). Lipids in bold contain a phosphate group.

| Lipid class | Fisher's P-value | Up or down-regulated* | Fold Change (low/high P) | T-Test P-value** |
| --- | --- | --- | --- | --- |
| **PE** | 0.00002 | Down | 0.6 | 0.006 |
| **PC** | 0.00010 | Down | 0.5 | 0.003 |
| **PA** | 0.00035 | Down | 0.5 | 0.017 |
| DG | 0.00053 | Down | 0.5 | 0.003 |
| **LPC** | 0.00069 | Down | 0.6 | 0.017 |
| **OxPG** | 0.00080 | Down | 0.5 | 0.011 |
| Cer-NS | 0.00443 | Down | 0.6 | 0.004 |
| DGTS | < 0.00001 | Up | 1.8 | 0.014 |
| TG | < 0.00001 | Up | 1.1 | 0.176 |
| DGDG | 0.00727 | Up | 1.2 | 0.204 |
| OxTG | 0.03723 | Up | 1.4 | 0.001 |

\*Up-regulated and down-regulated mean that the lipids were higher or lower, respectively, in leaves with low phosphorus content versus leaves with high phosphorus content based on fisher's exact test.
\*\*T-test (unequal variance, two-sided) of total intensities for sum of lipid class

Structures of certain upregulated and downregulated lipids are shown in Figure 2, showing structural similarity between PC and DGTS, for example. It is interesting to note that total DGTS had the greatest fold change increase in low P, as compared to other non-phosphorus containing membrane lipids, suggesting partial replacement of the dominant PC membrane lipid. Substitution of phospholipids by non-phosphate containing lipids was first reported in *Proteobacteria*^14^, where glycolipids replaced a large part of phospholipids in *Pseudomonas diminuta* so dramatically that in P-limited cultures, phosphate lipids were barely detectable (<0.3% of total polar lipids).

**Figure 2:**
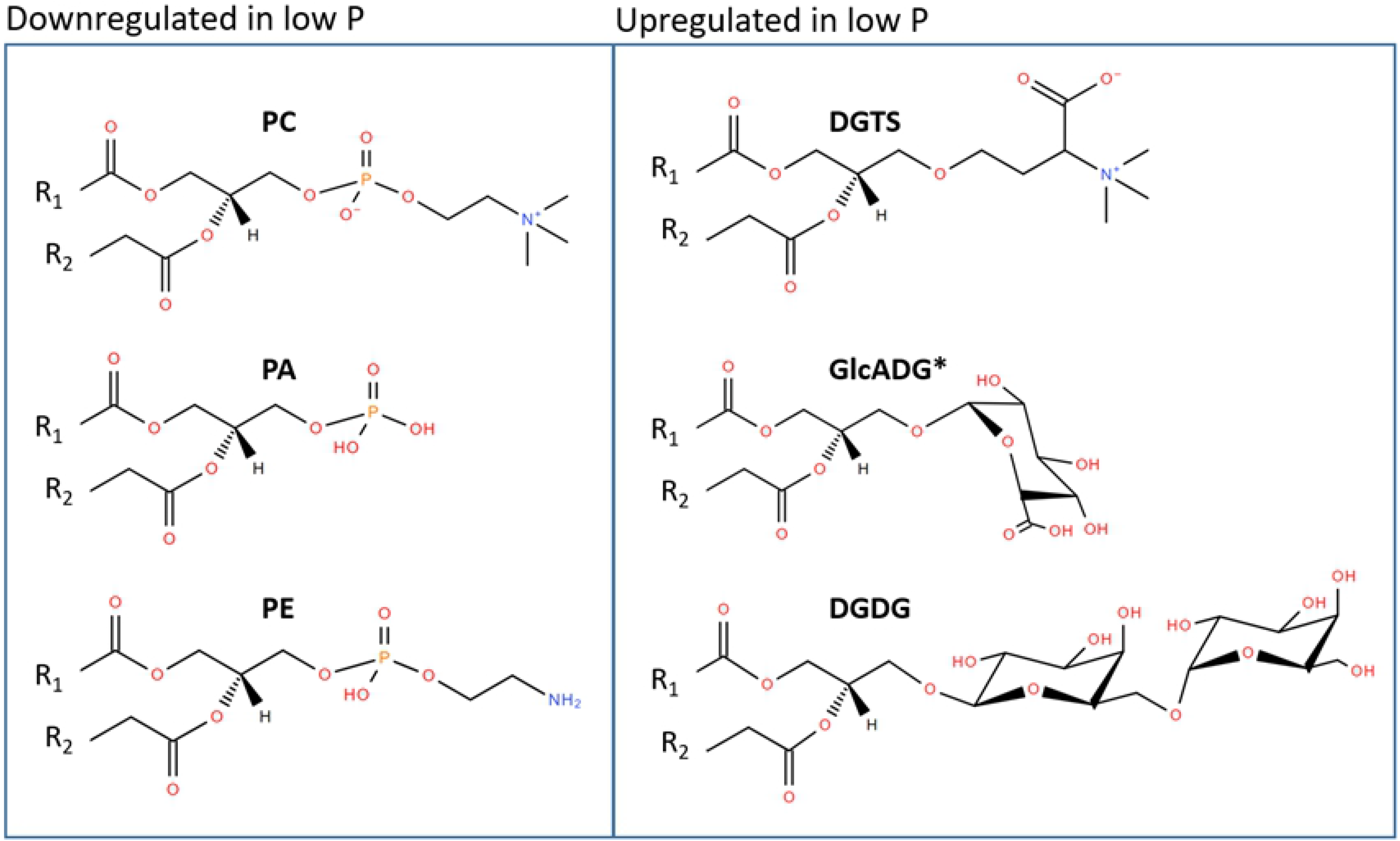
Examples of downregulated phospholipids that are key lipid species within the cell membrane, and significantly upregulated lipids that do not contain phosphorus. Lipid acronyms are defined as follows: phosphatidylcholine (PC), phosphatidic acid (PA), phosphatidylethanolamine (PE), diacylglyceryltrimethylhomoserine (DGTS), glucuronosyldiacylglycerols (GlcADG), and galactolipid digalactosyldiacylglycerol (DGDG). *for GlcADG only a few molecular lipid species were significantly upregulated

Since this landmark discovery, several lipid classes have been identified in a variety of diverse organisms to be involved in membrane lipid reconstructions during P starvation: SQDG was detected to substitute for phospholipids and thus to reduce P needs in *Arabidopsis* and certain species of picocyanobacteria.^15–17^ Similar modifications of membrane lipids, but with DGDG replacing phospholipids, have been reported in oat^18^ as well as in seven other species of monocots and dicots.^19^ DGTS (a P-free betaine-lipid analog of PC) has been reported to replace PC in fungi.^20^ So far, the only study revealing that membrane re-modeling is an important adaptation to low P concentrations in environmental mixed communities was reported for phytoplankton communities in the Sargasso Sea.^17^

Using LC-tandem MS and LipidMatch Flow software^19,20^ (methods in supplemental), we were able to identify that not all molecular species in a given lipid class showed the same trend and thus the data in Table 1 only shows a simplistic overview of the changes in lipid composition (SeaGrassData_Supplemental.xlsx and Figure S2). Under P-depleted conditions, the most significantly upregulated lipid species in *Thalassia* in terms of fold-change were actually GlcADG, SeaGrassData_Supplemental.xlsx, which were only recently discovered in the context of P starvation in *Arabidopsis*.^21^ Specifically GlcADG(16:0_18:2), fold change of 21, GlcADG(16:0_16:0), fold change of 7, and GlcADG (18:0_18:2), fold change of 7, were significantly higher under P-deplete conditions compared to high P (Figure S2). Interestingly, of the twelve GlcADG molecular species that were identified, only four were significantly upregulated (Hochberg corrected p-value < 0.05) and only the three listed above had fold changes above 2 (SeaGrassData_Supplemental.xlsx). Figure S1 shows examples of three GlcADG species identified by both MS-DIAL and LipidMatch, which had greatly differing fold changes. This impressively illustrates the use of MS and the urgent need for the identification of single molecular lipid species over other techniques that only analyze lipid classes (e.g. 2-D TLC) and explains why GlcADGs are not included in Table 1, which only shows overall changes in lipid classes.

Also, TGs were highly upregulated in P-deplete *Thalassia* leaves. TGs were also upregulated in nitrogen studies in the alga *Chlamydomonas reinhardtii*.^22^ In general in starvation conditions, membrane lipids are expected to decrease due to a shift towards TG synthesis^23^ as well as replacement by DGTS and DGDG^18^. We found that a significant number of DGDG species increased in P deficient seagrass leaves (Table 1, SeaGrassData_Supplemental.xlsx). Other lipids, which were downregulated under P-deplete conditions were diglycerides (DG) and ceramides (Cer-NS), (Table 1, SeaGrassData_Supplemental.xlsx). DGs are involved in DGTS, DGDG, and TG synthesis, all of which were upregulated in P-deficient *Thalassia* leaves. Still, more research is needed to understand the downregulation of DG and Cer-NS in P-deficient seagrass plants.

While the majority of the 32 SQDG species identified had fold changes greater than one (indicating upregulation; 27/32), only two were found to be significant (Hochberg corrected p-value < 0.05), namely SQDG (16:0_18:4) and SQDG (40:11), SeaGrassData_Supplemental.xlsx). Therefore, according to our study, SQDG had minor to no upregulation in concentration compared to TG, DGDG, DGTS, and certain GlcADG species and does not play an important role in remodeling of foliar membrane lipids under different P concentrations. While we cannot completely exclude that some of the 600 detected lipids originate from epiphytes or (endo)symbionts that were not completely removed by our washing steps, we are certain that the decrease in P-containing lipids reflect changes in the seagrass lipidome as phosphatidylcholine (PC), phosphatidic acid (PA), and phosphatidylethanolamine (PE) have previously been reported to be the main P-lipids in seagrasses.

In conclusion, we present evidence of a key cellular mechanism employed by a widely-distributed marine plant to thrive in nutrient-poor, oligotrophic conditions. These results not only explain the cellular mechanisms driving variability in turtlegrass P content, but also may potentially explain broader shifts in leaf structure or morphology under P-limitation, as membrane fluidity may be heavily influenced by lipid re-modelling. Understanding the biology of seagrasses and their adaptation to changing nutrient concentrations can help in conservation efforts. The lipid composition of seagrasses could be used as a biomarker to identify long-term nutrient limitation, which might not be detectable from periodic monitoring of nutrient concentrations in the surrounding waters.

## Authors’ contributions

JK analyzed the mass spec data, did the statistical analyses, and helped to write the manuscript, JC provided the samples and the nutrient data and edited the manuscript, JGC and LM performed the extractions and ran the mass spec, TG and US designed the study, JK and US wrote the manuscript. All authors read and approved the manuscript.

## Additional information

The authors declare that there are no financial and/or non-financial competing interests in relation to the work.

